# Impaired glutamate reuptake induces synaptic mistuning in rat hippocampal slices, that can be counteracted by ketamine

**DOI:** 10.1101/2022.01.25.477658

**Authors:** Erika Vazquez Juarez, Ipsit Srivastava, Maria Lindskog

## Abstract

Mistuning of synaptic transmission has been proposed to underlie many psychiatric disorders, with decreased reuptake of the excitatory neurotransmitter glutamate as one contributing factor. Synaptic tuning occurs through several diverging and converging forms of plasticity. By recording evoked field postsynaptic potentials in the CA1 area in hippocampal slices, we found that inhibiting glutamate transporters using DL-TBOA causes retuning of synaptic transmission, resulting in a new steady state with reduced synaptic strength and a lower threshold for inducing long-term synaptic potentiation (LTP). Moreover, we also found reduced threshold for LTP in a rat model of depression that has decreased levels of glutamate transporters. Most importantly, we found that the antidepressant ketamine counteracts the effects of increased glutamate on the various steps involved in synaptic retuning. We therefore propose that ketamine’s mechanism of action as an antidepressant is to restore adequate synaptic tuning.

## INTRODUCTION

A growing body of evidence indicates that impaired synaptic function is associated with a range of mental disorders (Frere and Slutsky, 2018; Vose and Stanton, 2017). A plethora of aspects of glutamate transmission and plasticity have lately been suggested to be affected in diseases that targets the brain, including enhanced glutamate transmission, a change in excitation/inhibition balance (Page and Coutellier, 2019), plasticity of excitatory synapses (Castrén and Antila, 2017) or a change in the homeostatic set-point (Frere and Slutsky, 2018). Further support for the relevance of excitatory synapse plasticity in psychiatric disorders in general—and depression in particular—comes from the finding that subanesthetic doses of the of *N*-methyl-D-aspartate (NMDA) receptor antagonist ketamine produce a fast-acting antidepressant effect (Berman et al., 2000; Zarate et al., 2006). Although our understanding of ketamine’s mechanism of action is still ambiguous (Duman et al., 2019; Wei et al., 2021), ketamine has been shown to beneficially affect glutamatergic transmission (Abdallah et al., 2015; Kavalali and Monteggia, 2020). The therapeutic effects were originally attributed to a reduction in excitatory activity by inhibiting NMDA receptors; however, further research revealed that ketamine can have complex effects on the balance between excitation and inhibition and on synaptic plasticity (Cornwell et al., 2012; Duman et al., 2019; Moda-Sava et al., 2019), including——paradoxically—an increase in excitatory transmission through decreased inhibition and/or synaptic plasticity mechanisms (Zanos and Gould, 2018).

The evidence for ketamine, and other psychiatric drugs, affecting glutamate transmission reinforces the idea that to ensure healthy brain function, a balanced excitatory synaptic transmission is needed. To ensure plasticity while keeping the activity within a healthy range, various forms of plasticity interact. For example, in Hebbian (i.e., activity-dependent) plasticity, coordinated neuronal activity generates a positive feedback process that reinforces the strength of specific synapses with high levels of activity (Nicoll, 2017). Conversely, homeostatic plasticity uses negative feedback to downregulate synaptic activity in response to high levels of global activity (Turrigiano, 2012). The interplay between these two forms of plasticity is fundamental for ensuring balanced neuronal activity while providing the flexibility needed to maintain brain function (see for example (Keck et al., 2017; Stampanoni Bassi et al., 2019). However, our knowledge regarding the interaction between various forms of plasticity is based to a large extent on indirect measurements such as changes in receptor expression and synapse morphology (Piva et al., 2021; Thiagarajan et al., 2006).

Interestingly, one key mechanisms for keeping excitatory neurotransmission balanced - namely glutamate reuptake - appears to be affected in many mental disorders. In particular, the astrocytic glutamate transporters EAAT-1 and EAAT-2, which take up the majority of synaptic glutamate (Murphy-Royal et al., 2017), have consistently been reported to be expressed at reduced levels in individuals with cognitive and/or psychiatric disorders (Parkin et al., 2018; Rajkowska and Stockmeier, 2013). Increased levels of extracellular glutamate are typically believed to lead to excitotoxicity; however, reduced glutamate reuptake can induce a number of adaptive changes and does not necessarily induce cell death (Barnes et al., 2020; Gomez-Galan et al., 2013). To date, strikingly little is known regarding the process by which synaptic tuning is affected when glutamate transporter levels are reduced.

Considering the consistent finding of downregulated glutamate transporters both in animal models of depression and in patients with depression, and given the association between synaptic plasticity and depression, it is of great relevance to understand how the tuning of excitatory synapses is affected by reduced glutamate reuptake. We therefore measured evoked synaptic responses in rat hippocampal slices while blocking glutamate transporters and found that a series of retuning steps lead to a new steady state in which the threshold for inducing plasticity is reduced. Mover, we found that ketamine was effective at reversing all of these retuning steps, providing new insights into its mechanism of action as an antidepressant.

## RESULTS

### Blocking glutamate transporters induces synaptic retuning and increased levels of extrasynaptic glutamate

The acute effect on reduced glutamate transporter function was examined by recording field excitatory postsynaptic potentials (fEPSPs) in the CA1 area in response to presynaptic activation of the Schaffer collaterals. Bath application of the glutamate transporter inhibitor DL-TBOA (DL-threo-beta-benzyloxyaspartate) caused a gradual but transient decrease in the magnitude of the fEPSP response, quantified by the slope of the rising phase of the evoked fEPSPs (see Methods; Fig. 1A). In the continued presence of DL-TBOA, the magnitude of the response decreased with an average rate of -3.1±0.3% per min (Fig. 1B), with complete loss of the response within 40 minutes (Fig. 1A); this was followed by recovery of the response to approximately 44% of baseline (Fig. 1A,B). In contrast, DL-TBOA did not affect fiber volley amplitude (data not shown), indicating that the stimulating input was unchanged. Interestingly, washing out DL-TBOA increased fEPSP magnitude further, albeit to only approximately 56% of baseline, this reflecting a new steady state. Similar results were obtained using TFB-TBOA, another glutamate transporter inhibitor with high specificity for transporters expressed in astrocytes (Suppl. Fig. S1A); however, because the effects of TFB-TBOA were more variable, DL-TBOA was used for subsequent experiments.

**Fig. 1:**
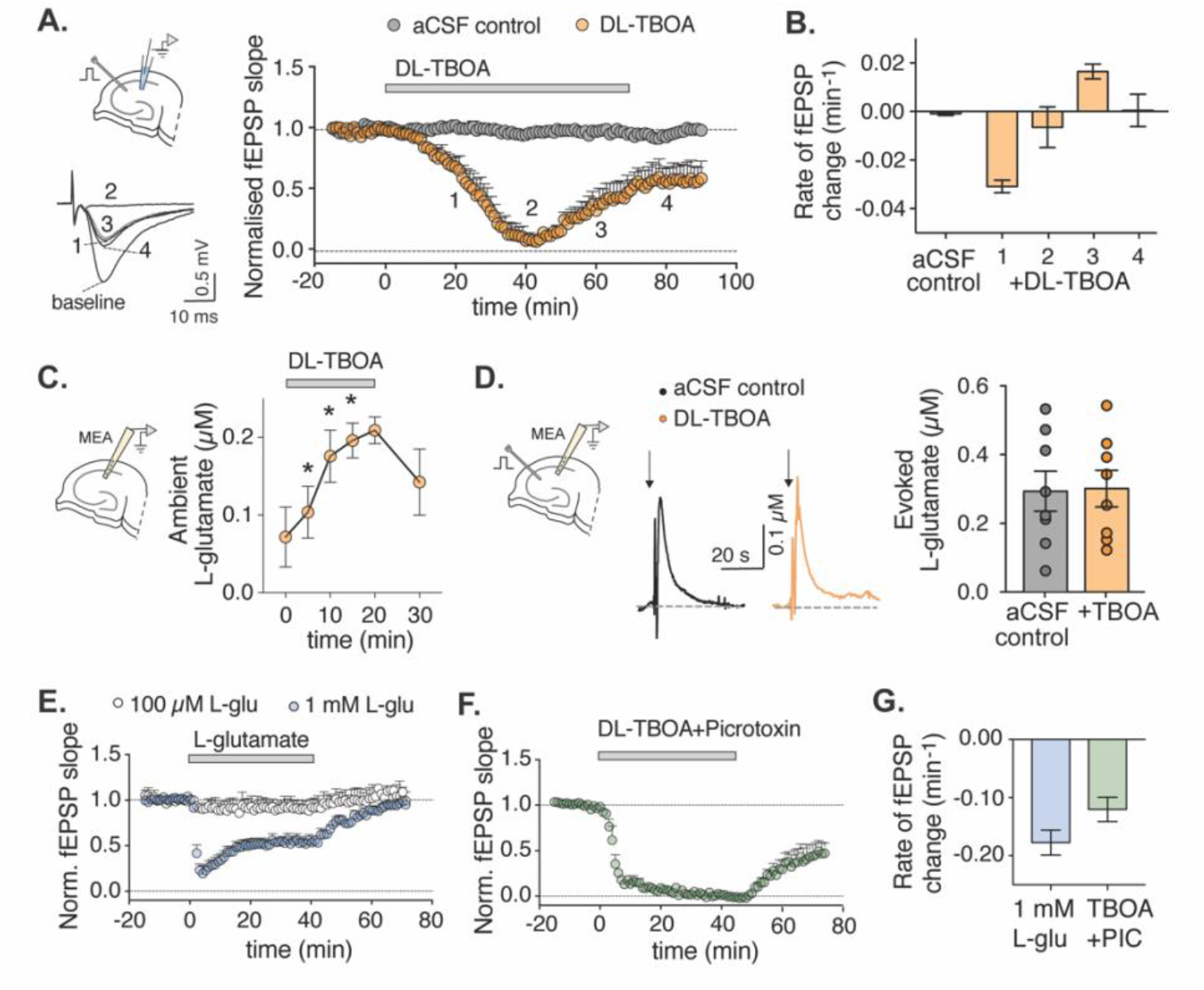
Increasing extrasynaptic glutamate by blocking glutamate transporters induces synaptic retuning. A) Blocking glutamate transporters induces a dynamic change over time of fEPSP magnitude in rat hippocampal slices, shown as normalized fEPSP slope. The Schaffer collaterals were stimulated at 0.1 Hz throughout the experiment while recording from the stratum radiatum at CA1 (scheme at left). Where indicated, the glutamate transporter inhibitor DL-TBOA (50 µM) was applied and then washed out (n=4 and 5 slices for aCSF control and DL-TBOA, respectively). The fEPSP slope over time is plotted as average of 6 stimulation pulses per minute in this and subsequent figures. One-minute averaged example traces (inset bottom left) of fEPSP recorded at baseline and at indicated time-points (20, 40, 60 and 80 minutes after DL-TBOA application). B) Rate of change in the fEPSP response at the indicated time periods, measured as the slope of a first order linear model fitted to the fEPSP time-response. C) Extracellular glutamate recorded in real-time with an enzyme-coated microelectrode before and after application of 50 µM DL-TBOA (n=11 slices). F) Example traces (left) of extracellular glutamate recordings during Schaffer collateral stimulation (arrows) and average peak glutamate concentration (right); n=7 and 8 slices for control and 50 µM DL-TBOA, respectively. E) 1 mM, but not 100 μM, L-glutamate induced similar retuning of synaptic transmission as blocking glutamate transporters. Glutamate was applied where indicated (n=5 slices each). F) The GABA receptor antagonist picrotoxin (50 µM) accelerated the DL-TBOA–induced retuning when added together with DL-TBOA. G) Summary of the rate of change in the fEPSP response measured 1-5 min after drug application (indicated by the gray shaded area in panels E and F), measured as the slope of a first order linear model fitted to the fEPSP time-response. In this and subsequent figures, error bars denote the standard error of the mean (SEM). *p<0.05 (ANOVA followed by post-hoc test).

Importantly, we ruled out a toxic effect of DL-TBOA in the slices by propidium iodide uptake experiments (a measure of cell death), finding no significant difference between DL-TBOA– treated slices and control-treated slices (Suppl. Fig. S1B). Next, direct recordings of extracellular levels of glutamate using enzyme-based amperometric glutamate sensors, showed that DL-TBOA significantly increased the basal level of extracellular glutamate from approximately 80 nM to approximately 200 nM, even in the absence of external stimulation (Fig. 1C). Interestingly, however, DL-TBOA did not affect peak glutamate levels after synaptic activation, as stimulation of the Schaffer collaterals increased extracellular glutamate to the same level (approximately 300 nM) regardless of the presence or absence of DL-TBOA (Fig. 1D).

The relatively rapid increase in extracellular glutamate induced by DL-TBOA compared to the slower, delayed effect of DL-TBOA on fEPSP amplitude (compare Fig. 1C with Fig. 1A) suggests that the neuronal network can initially compensate for the effects of an increase in extracellular glutamate, although this compensatory mechanism is likely overwhelmed when extracellular glutamate is chronically increased. Consistent with this suggested resilience of the system, bath application of a relatively low concentration of glutamate (100 µM) had no significant effect on fEPSPs (Fig. 1E), whereas application of 1 mM glutamate induced a change in fEPSPs that was qualitatively similar to the effects observed with DL-TBOA (Fig. 1E,G).

One possible mechanism for the decrease in the excitatory synaptic response that we observe after prolonged increase in extracellular glutamate is an increased inhibition via activation of GABAergic neurons. However, co-application of the GABA receptor inhibitor picrotoxin together with DL-TBOA caused a rapid, sustained decrease in fEPSP magnitude, with an average rate of change of -12±2.1% per min (Fig. 1F,G). Thus, blocking GABA receptors accelerated the DL-TBOA–mediated decrease in fEPSP magnitude. Another mechanism of fEPSP reduction in response to increased extracellular glutamate would be AMPA receptor desensitization. Surprisingly, we found that cyclothiazide, a compound that prevents AMPA receptor desensitization, rapidly decreased fEPSP magnitude (Suppl. Fig. S1C,D). This unexpected result could be explained by the fact that cyclothiazide also affects GABA_A_ receptors(Deng and Chen, 2003), a notion supported by the remarkably similar results obtained between cyclothiazide and picrotoxin.

### The decrease in fEPSP magnitude after blocking glutamate transporters is mediated by NMDA receptors

Having ruled out the effect of inhibition in the reduction of fEPSP magnitude, and given that NMDA receptors play a key role in synaptic plasticity, we examined the effect of various concentrations of NMDA on evoked fEPSP in slices. Similar to the results obtained with glutamate, we found that NMDA decreased the magnitude of evoked fEPSPs in a concentration-dependent manner; specifically, 1 µM NMDA had no effect, while 5, 7, and 10 µM NMDA decreased fEPSP magnitude by approximately 37%, 64%, and 100%, respectively, with 10 µM NMDA reaching its maximum effect within 4 minutes (Fig. 2A,B). Interestingly, the effects of both 5 and 7 µM NMDA were fully reversed after wash-out, with fEPSP magnitude returning to baseline levels; in contrast, 10 µM NMDA caused a long-lasting change in synaptic strength that remained after wash-out (Fig. 2A). Moreover, pretreating slices with either the competitive NMDA receptor antagonist AP5 or the NMDA receptor pore blocker MK-801 prevented the of NMDA receptors was verified by recording fEPSPs in the presence of 25 µM NBQX to block AMPA receptors and 0.2 mM MgCl_2_ to favor NMDA receptor opening; under these conditions, we observed a transient increase in the magnitude of the NMDA receptor-mediated component of the fEPSP upon addition of DL-TBOA (Fig. 2E,F and Suppl. Fig. S2B).

**Fig. 2:**
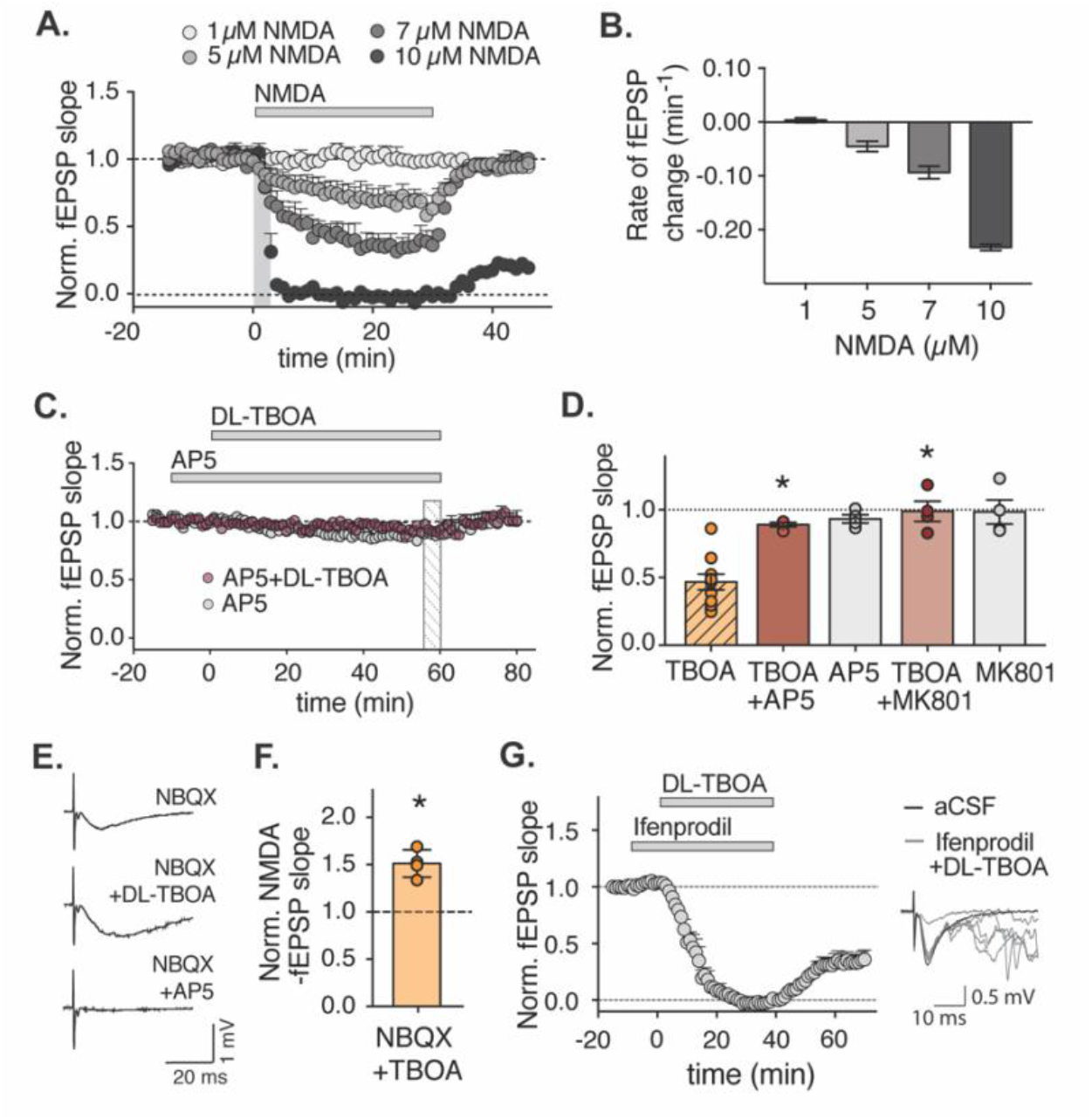
The decrease in fEPSP magnitude requires NMDA receptor activity. A) Synaptic retuning can be mimicked by NMDA in a concentration-dependent manner. Time course of normalized fEPSP slope; where the indicated concentrations of NMDA were added as shown (n=3-5 slices each). B) Rate of change of the fEPSP measured as the slope of a first order linear model fitted to the fEPSP time-response during the first 4 min after NMDA application. C) Synaptic retuning is blocked by 10 µM AP5. D) Average normalized fEPSP slope measured 56-60 minutes after application of the indicated compounds (shaded area in C), hatched bar indicates that the data has been shown in previous figure; *p<0.05 vs. TBOA (ANOVA). E) NMDA synaptic responses are initially increased when glutamate transporters are blocked. Example traces of NMDA receptor–mediated field potentials recorded in the presence of 25 µM NBQX and 0.2 mM MgCl_2_; bath application of 50 µM AP5 eliminated the response (bottom trace). F) Average fEPSP slope measured 5 min after application of NBQX + DL-TBOA, normalized to NBQX alone (n=4 slices each). *p<0.05 (Mann-Whitney U test). G) Normalized fEPSP slope; where indicated, 10 µM ifenprodil and 50 µM DL-TBOA were applied (n=5 slices each). Shown at the right are exemplar traces of fEPSPs, showing epileptiform activity 10 min after the addition of DL-TBOA in the presence of ifenprodil (gray traces) compared to baseline (black trace).

Increased extracellular glutamate due to reduced glutamate reuptake could lead to increased activation of extrasynaptic NMDA receptors, which are described to contain a higher proportion of the GluN2B subunit (Papouin et al., 2012). To test this possibility, we used the GluN2B subunit–specific antagonist ifenprodil. Surprisingly, we found that the presence of 10 µM ifenprodil had no effect on the DL-TBOA–induced decrease in fEPSP magnitude, but induced an epileptiform activity pattern (Fig. 2G and Suppl. 2C). This finding suggests that GluN2B subunit–containing NMDA receptors are indeed activated to a higher extent when glutamate transporters are blocked, and activates responses to downregulate neuronal activity via a mechanism independent of the reduction in synaptic response.

To examine the possibility that NMDA receptor activation triggers internalization of AMPA receptors, DL-TBOA treated slices were incubated with biotin to assess the amount of AMPA receptors present on the surface of the membrane. Indeed, the amount of biotinylated AMPA was reduced in DL-TBOA treated slices compared to control (Suppl. Fig. 3, A-C), however, not to a level that alone could explain the strong reduction in fEPSP magnitude that we observed. Thus, we also examined the possibility that increased extrasynaptic glutamate reduces presynaptic release through activation of presynaptic metabotropic glutamate receptors (mGluR2/3) with the mGluR2/3 antagonist LY341495 (1 µM). This treatment induced a slight reduction in the rate of decrease of the fEPSP magnitude (Suppl. Fig 3, D-E) indicating that the reduction in fEPSP magnitude that we observe is a consequence of at least two interacting mechanisms.

### The recovery of fEPSP magnitude is dependent on presynaptic activity

Unlike the decrease in fEPSP magnitude, recovery of the evoked response did not require NMDA receptor activation (Fig. 3A). In contrast, the recovery—but not the initial decrease—was dependent on the frequency of the stimulation used to evoke field responses (Fig. 3B,C). When fEPSPs were evoked every 15 minutes (i.e., at approximately 0.001 Hz), we found no recovery of the evoked fEPSPs; upon switching the stimulation frequency to 0.1 Hz, however, the fEPSP response recovered, indicating that recovery from the non-responding state in the presence of DL-TBOA requires presynaptic activation (Fig. 3C). Interestingly, when fEPSPs were evoked at the slower frequency (i.e., at approximately 0.001 Hz), the DL-TBOA–induced decrease in magnitude occurred at a considerably slower rate compared to previous experiments in which the stimuli were applied at 0.1 Hz (see Fig. 3D).

**Fig. 3:**
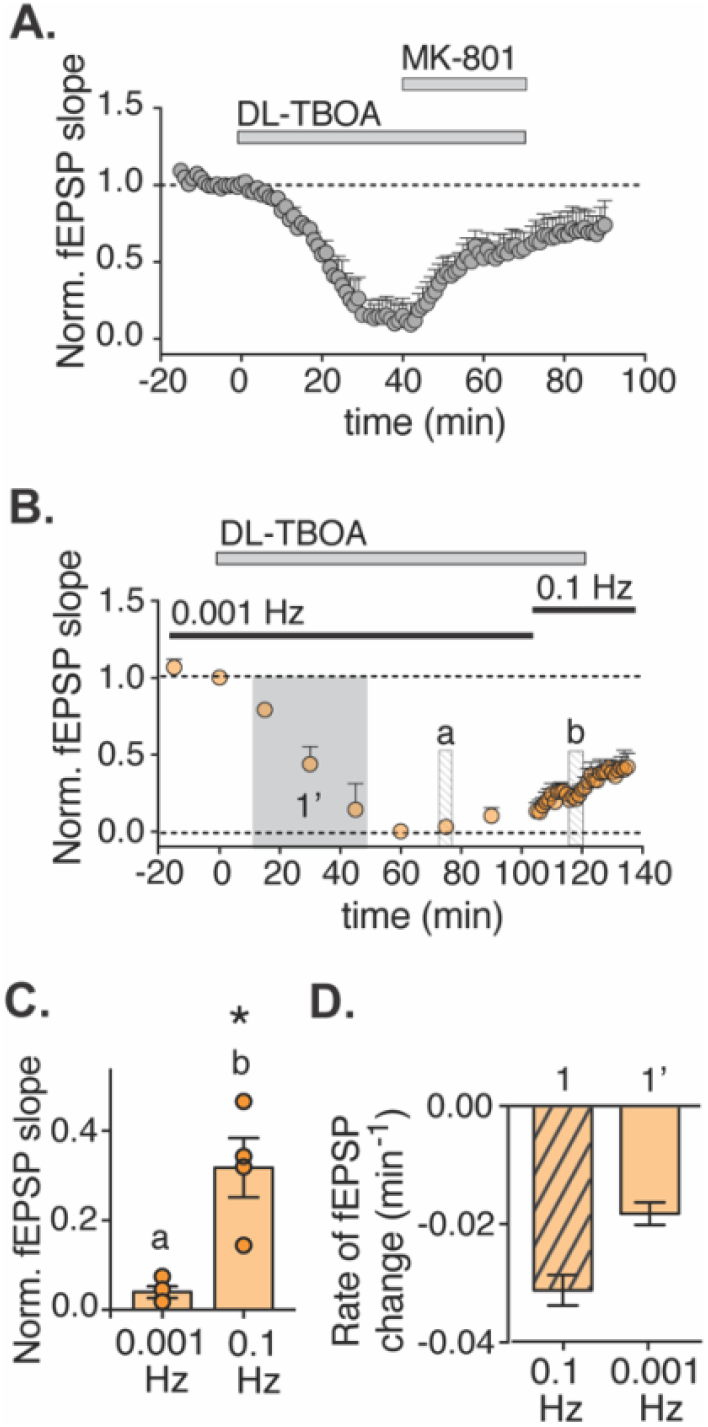
The recovery of fEPSPs is dependent on stimulation frequency but does not require NMDA receptor activity. A) Time course of normalized fEPSP slope; where indicated, DL-TBOA and MK-801 were added (n=4 slices). B) Time course of normalized fEPSP slope measured using the indicated stimulation frequencies; where indicated, DL-TBOA was added (n=5 slices). C) Summary of normalized fEPSP slope measured at the indicated times in panel B. *p<0.05 (ANOVA). D) Summary of the rate of change of the fEPSP at the indicated times in Fig. 1A (“1”) and panel B (“1’”); n=4-5 slices each; hatched bar indicates that the data has been shown in previous figure.

### The threshold for synaptic potentiation is decreased following retuning

Our finding that fEPSP magnitude does not return to baseline levels after its initial decrease and partial recovery in the presence of DL-TBOA during 0.1 Hz stimulation (see Fig. 1A) suggests that the synapses have entered a new steady state that prevailed even after wash-out of DL-TBOA. To examine the synaptic properties under this putative new steady state, we recorded miniature excitatory postsynaptic currents (mEPSCs) in slices in which the synapses had been retuned with DL-TBOA and in control slices in which DL-TBOA was replaced with aCSF (Fig. 4A,B). We found no significant difference in either the frequency or amplitude of mEPSCs between slices that had been retuned with DL-TBOA compared to control slices (Fig. 4C,D), although there was a trend to a decrease in frequency. This could be an indication of a presynaptic change, since a change in mEPSC is typically associated with a decrease in presynaptic probability of release. To examine this further we analyzed the paired-pulsed facilitation (PPF) in slices that had undergone the retuning protocol versus control treated slices and observed a significant increase in PPF in retuned slices (Fig. 4E), again suggesting a decreased presynaptic release probability. We then examined the ability to induce long-term potentiation (LTP) by applying two trains of 10 stimuli at 5 Hz (i.e., θ-burst stimuli) and found no difference in LTP between DL-TBOA–treated slices and control slices (Fig. 4F). To our surprise, however, we found that a subthreshold θ-burst stimulation elicited only short-term potentiation in control slices but induced significant LTP in slices that had been retuned with DL-TBOA (Fig. 4G); this suggests that the new steady state induced by increasing extracellular glutamate renders the synapses more susceptible to long-term plasticity.

**Fig. 4:**
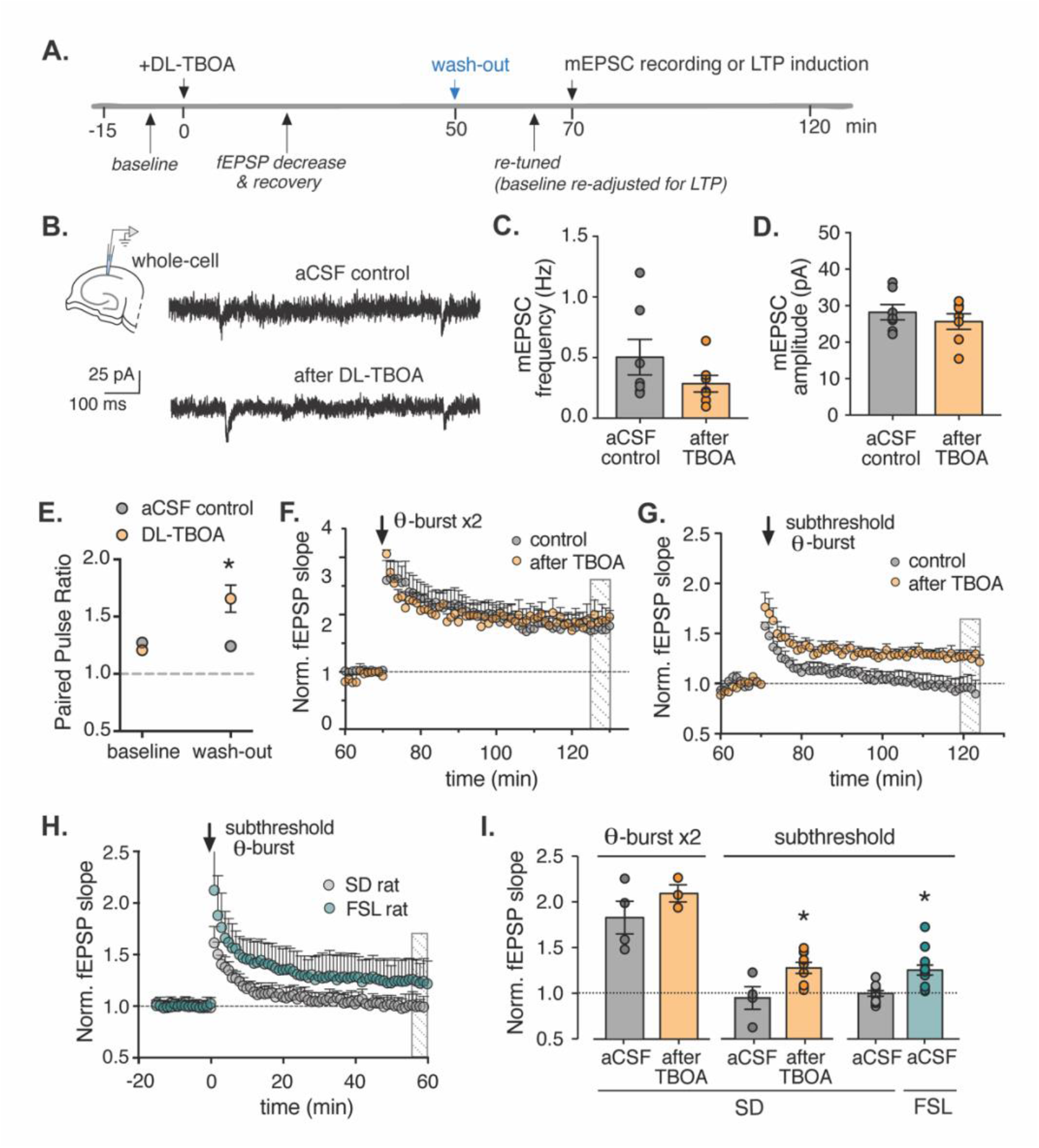
Decreasing glutamate uptake lowers the threshold for inducing long-term potentiation. A) Protocol used to induce a new steady state in hippocampal slices by applying 50 µM DL-TBOA for 40 minutes while stimulating the Schaffer collaterals at 0.1 Hz. Control slices are treated similarly but are perfused with aCSF instead of DL-TBOA. B) Representative traces of patch-clamp recordings in control and retuned slices; tetrodotoxin was present throughout the recordings to isolate mEPSCs. C-D) Summary of the frequency (C) and amplitude (D) of mEPSCs recorded in control slices and retuned slices (n=7 slices each). E) Paired-pulse facilitation ratio (PPR) obtained by dividing the fEPSP slope evoked by the second pulse (50 ms inter-stimulus interval) by the fEPSP slope evoked by the first pulse; PPR in control (n=3) and DL-TBOA-treated slices (n=6), was calculated at the baseline period and after wash-out indicated in the time-line showed in A. F) Time course of normalized fEPSP slope in control slices (n=4) and retuned slices (n=3); where indicated, a standard LTP-inducing theta-burst (θ-burst) was applied. G) Time course of normalized fEPSP slope in control slices (n=4) and retuned slices (n=10); where indicated, a subthreshold θ-burst stimulation was applied. H) A subthreshold θ-burst stimulation was applied in slices obtained from either Sprague-Dawley (SD) rats FSL rats. I) Summary of normalized fEPSP slope averaged over 5 min in the shaded regions in in panels E, F, and G. *p<0.05 versus the corresponding baseline (ANOVA).

The finding that an acute inhibition of glutamate transporters with DL-TBOA induces a new steady state with a reduced threshold for LTP prompted us to investigate the translational relevance of this phenomenon. We therefore examined the effect of subthreshold stimulation in hippocampal slices prepared from FSL (Flinders Sensitive Line) rats, an established model of depression in which glial glutamate transporters are downregulated (Gomez-Galan et al., 2013). We found that the same subthreshold θ-burst stimulation that did not induce LTP in slices prepared from control (Sprague-Dawley, or SD) rats, was sufficient to induce robust LTP in slices prepared from FSL rats (Fig. 4H), with results similar to DL-TBOA–treated slices prepared from SD rats (Fig 4I).

### The antidepressant drug ketamine affects the retuning and new steady state at several steps

Given that the fast-acting antidepressant drug ketamine is an NMDA receptor antagonist, we tested whether a clinically relevant dose of ketamine can prevent the DL-TBOA–induced decrease in fEPSP magnitude. Similar to our results obtained with the NMDA receptor blocker MK-801 and the NMDA receptor antagonist AP-5, we found that 20 µM ketamine eliminated the ability of DL-TBOA to decrease fEPSP magnitude in rat hippocampal slices (Fig. 5A,B).

**Fig. 5:**
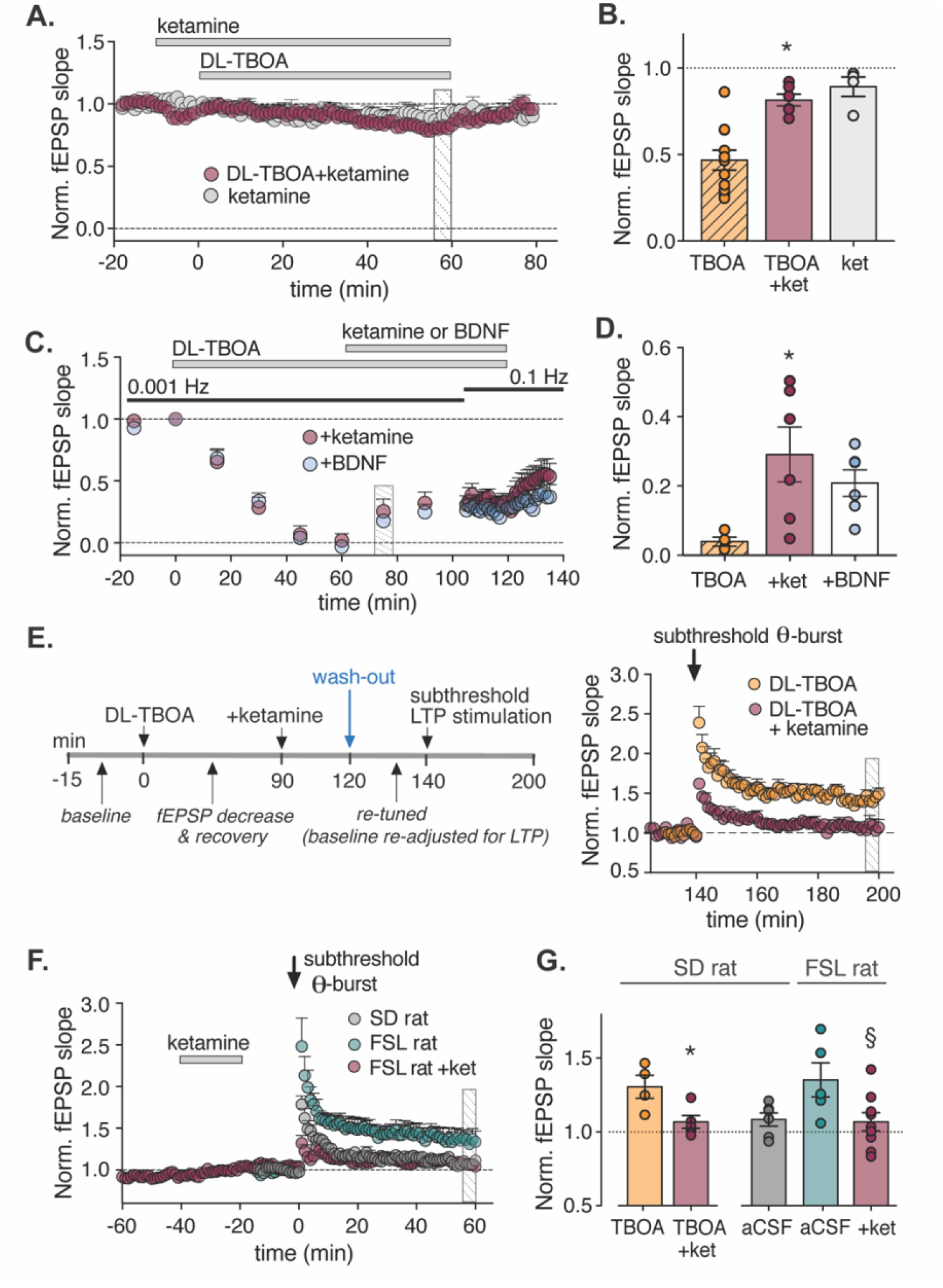
Ketamine affects synaptic tuning at multiple steps. A) Time course of normalized fEPSP slope in acute slices from SD rats; 20 µM ketamine and/or DL-TBOA were applied as indicated (n=4-6 slices each). B) Average fEPSP slope normalized to baseline measured 55-60 minutes after the application of 50 µM DL-TBOA (data from Fig. 1A; hatched bar indicates that the data has been shown in previous figure), 20 µM ketamine (n=4 slices), or 20 µM ketamine + 50 µM DL-TBOA (n=6 slices). C) Time course of normalized fEPSP slope while applying stimuli at the indicated frequencies; where indicated, DL-TBOA and either ketamine or BDNF were applied (n=6 slices each). D) Average fEPSP slope measured 15 minutes after ketamine or BDNF (see panel B) versus DL-TBOA alone (from Fig. 1). *p<0.05 versus DL-TBOA alone (ANOVA). E) Left: Protocol used to test the effect of pretreating slices with 20 µM ketamine on synaptic plasticity in hippocampal slices from SD rats. Right: Time course or re-normalized fEPSP slope; where indicated, a subthreshold θ-burst stimulation was applied (n=4-5 slices each). F) Time course of normalized fEPSP slope measured in slices from SD and FSL rats. Where indicated, ketamine was applied to the FSL slices (n=10-13 slices each). G) Summary of normalized fEPSP slope measured 1 h after LTP induction under the indicated conditions *p<0.05 versus DL-TBOA and §p<0.05 versus the corresponding aCSF group (Student’s t-test).

A previously suggested mechanism of action for ketamine is suppression of spontaneous NMDA receptor activation, inducing a rapid increase in evoked transmission via the BDNF-dependent recruitment of AMPA receptors (Nosyreva et al., 2013). We therefore tested whether ketamine can affect the recovery phase of synaptic tuning measured 60 minutes after application of DL-TBOA under low-frequency (i.e., 0.001 Hz) stimulation (Fig. 5C). We found that the addition of either 20 µM ketamine or 10 ng/ml BDNF caused a significant recovery of fEPSP magnitude (Fig. 5C,D).

Finally, we examined whether briefly treating slices in the new steady state with ketamine could reverse the increased susceptibility to LTP induced with subthreshold θ-burst stimulation. We therefore treated SD slices prepared with DL-TBOA during 0.1-Hz stimulation to induce the new steady state, followed by treatment with ketamine for 30 minutes; after a wash-out period of 20 minutes, we then applied subthreshold θ-burst stimulation. We found that subthreshold stimulation failed to induce LTP in the ketamine-treated slices (Fig. 5E). Moreover, treating slices obtained from FSL rats with ketamine also eliminated LTP induced by subthreshold stimulation (Fig. 5F,G). Thus, brief application of a low dose of ketamine prevented all the retuning steps that we have observed in reponse to blocking glutamate uptake.

## DISCUSSION

Here, we report that inducing synaptic retuning by altering synaptic glutamate levels causes both a quantitative change in synaptic strength and a change in the susceptibility to induce long-term plasticity. Thus, in contrast to the often-used metaphor of the synapse serving as a “volume control” in which neurotransmission can be dialed up or down, synaptic tuning may actually be more similar to a gearbox, in which the overall drive and acceleration are modulated. The ability to fine-tune synaptic transmission through interacting homeostatic and feed-forward mechanisms would be at the core of achieving a robust—yet highly plastic—system.

In our experiments, the time course of the first retuning step—the reduction in fEPSP magnitude—as well as the dependence of this process on NMDA receptors, is similar to what is described for long-term depression (LTD) (Collingridge et al., 2010) in contrast to homeostatic scaling where the downregulation is typically lower and slower (Ibata et al., 2008). Consistent with recent work (Barnes et al., 2020), we rule out any simple explanation of this downregulation, since neither GABA inhibition (Fig. 1F), blocking AMPA receptor desensitization (Suppl. Fig. S1C,D) or blocking GluN2B (Fig. 2G) prevents the initial step of the re-tuning. These results are consistent with previous studies showing that blocking specific intracellular pathways can prevent DL-TBOA–mediated effects on synaptic transmission independently (Barnes et al., 2020; Falcón-Moya et al., 2020). Furthermore, blocking mGluR2/3 receptors that are known as presynaptic auto-receptors, delays the effect of DL-TBOA, and in addition we see a slight decrease in membranous AMPA receptors, that we interpret as a postsynaptic adjustment, after DL-TBOA treatment (Suppl. Fig. 3). Taken together, our results indicate that the synaptic retuning taking place upon increased extracellular glutamate levels is transsynaptic – involving pre as well as postsynaptic changes.

NMDA receptors have an established role in plasticity; in particular, extrasynaptic receptors are described to play an essential role in maintaining synaptic tuning (Papouin and Oliet, 2014). Here, we show that NMDA receptor activation is required for the initial reduction in synaptic strength; moreover, the rate of retuning increases with increased concentrations of NMDA. Interestingly, however, we also found that the rate of retuning is increased by blocking GABA receptors with picrotoxin and decreased when activity was driven using low-frequency stimulation. Thus, retuning seem to be an effect of general postsynaptic depolarization together with NMDA receptor activation. This is in good agreement with what has been described in the visual cortex, where monocular deprivation induces LTD through noncoherent input and extended postsynaptic depolarization (Linden et al., 2009). Moreover, in the visual cortex, prior synaptic activity and NMDA receptor activation reduces the threshold for plasticity (Cooper and Bear, 2012; Rodriguez et al., 2019), similar to our findings in the hippocampus. Due to its relatively easy access, the visual cortex has served as a model for studying plasticity and adaptation in response to changes in activity (Kaneko and Stryker, 2017), and our results suggest that reducing glutamate uptake in the hippocampus and monocular deprivation in the visual cortex use similar retuning mechanisms. Hence, our knowledge regarding the way in which synaptic tuning mechanisms in the visual cortex adjust to global changes in activity may also be applicable to other cortical regions, including the hippocampus.

An intriguing finding in this work is that although we can record no synaptic response after 40 minutes of DL-TBOA, the subsequent recovery is dependent of presynaptic stimulation (Fig. 4). In the case of inactivity-induced homeostatic scaling, the underlying mechanism involves the spontaneous release of glutamate driving phosphorylation of eEF2 (eukaryotic elongation factor 2) and inhibition of local translation (Autry et al., 2011; Sutton et al., 2007). Thus, in our case it is reasonable to speculate that presynaptic activity in the context of inhibited glutamate transporters relieves this effect on translation. Consistent with this notion, in the absence of presynaptic activity the recovery of fEPSPs can be induced by BDNF or ketamine (Fig. 5C,D), which is supported by previous findings (Kavalali and Monteggia, 2020).

The existence and robustness of a homeostatic “set point” of activity may be a fundamental feature of neuronal circuits achieved by a wide range of interacting mechanisms. Such an activity set point can ensure that brain activity is balanced despite perturbations, changes in activity levels over time, and even injury (Ma et al., 2019; Turrigiano, 2017). However, the price to pay for this stable system is that changes in the set point itself—as suggested in certain psychiatric disorders such as depression (Vose and Stanton, 2017) and Alzheimer’s disease (Styr and Slutsky, 2018)—can be devastating. The set point of neuronal activity is regulated by synaptic plasticity/homeostasis (Turrigiano, 2012), excitatory/inhibitory balance (Field et al., 2020), and cellular excitability (Styr and Slutsky, 2018). Here, we found that increasing extracellular glutamate by blocking glutamate transporters induces a long-lasting downregulation of the set point that remains even after the glutamate transporters are no longer blocked. Thus, blocking glutamate transporters induces a long-lasting change in steady-state synaptic activity and reduces the threshold for plasticity. Although in this case we change the set-point to a pathological state, the opposite manipulation could be tested for therapeutic effects. Moreover, the fact that the majority of glutamate transporters are expressed on astrocytes underscores the importance of these cells in maintaining balanced neuronal activity, as well as their potential as a therapeutic target for the treatment of certain psychiatric and cognitive disorders (Escartin et al., 2021). Interestingly, glutamate transporter localization is dynamic and dependent on astrocyte morphology and membrane turn-over (Michaluk et al., 2021) thus differences in astrocyte morphology that are well described in different disease models may have functional consequences for glutamate uptake (Escartin et al., 2021).

This study was designed to investigate the consequences of reduced glutamate transport on synaptic activity, given the pathogenic role that this reduction appears to play in a wide range of psychiatric disorders, including major depression (Parkin et al., 2018). Indeed, we found that the threshold for inducing LTP was significantly reduced in hippocampal slices prepared from FSL rats, an established animal model of depression with reduced expression of glutamate transporters (Gomez-Galan et al., 2013). We have previously reported that LTP amplitude is reduced in the FSL rat (Gomez-Galan et al., 2013), and taken together, the reduced threshold and amplitude, could lead to an unstable network with increased randomness, a notion consistent with EEG recordings of patients with depression (Zhang et al., 2018). Interestingly, our results indicate that the NMDA receptor antagonist ketamine—used clinically as a fast-acting antidepressant—affects the “synaptic gearbox” at several sites. Not surprisingly, we found that ketamine prevents the decrease in synaptic strength induced by DL-TBOA, mimicking the effects of other NMDA receptor antagonists. As the initial decrease is independent of the GluN2B receptor, it is unlikely that ketamine acts onlythrough extrasynaptic receptors as has been proposed (Miller et al., 2014). Furthermore, ketamine was also effective when applied in the presence of DL-TBOA during low-frequency (0.001-Hz) stimulation, and ketamine prevented the increased susceptibility to LTP both in DL-TBOA–treated slices obtained from wild-type rats and in slices obtained from FSL rats. Thus, ketamine appears to stabilize synaptic tuning regardless of the synapse’s previous state. In this respect, it is interesting to note that structural plasticity was recently reported to be required the ketamine’s sustained effects, but not for the induction (Moda-Sava et al., 2019). Thus, our finding of synaptic retuning suggests a potential mechanism underlying the rapid initial response to ketamine.

Taken together, these results support the emerging hypothesis that depression may be explained as a dysregulation of synaptic tuning rather than a change in excitation or inhibition *per se*. The interaction of different plastic mechanisms and the resulting synaptic retuning may therefore represent a promising new framework for developing new treatments for depression and related psychiatric disorders.

## MATERIALS AND METHODS

### Animals

All experiments were approved by the regional committee for animal research of Stockholm North in Sweden (N13/15). Unless stated otherwise, male adult (6-9 weeks of age) rats were used. Sprague-Dawley (SD) rats were supplied from Janvier Laboratories, and FSL rats were bred at the Karolinska Institute. All rats were group-housed at a 12-h light/dark cycle in the animal facility at Karolinska Institute and had *ad libitum* access to food and water. For slice preparation, the rats were deeply anesthetized with isoflurane and decapitated shortly after loss of the corneal reflex.

### Drugs

DL-TBOA, TFB-TBOA, DL-AP-5 sodium salt, MK-801 maleate, ketamine hydrochloride, picrotoxin, cyclothiazide, NBQX disodium salt, ifenprodil hemitartrate, LY341495 and tetrodotoxin were purchased from Tocris Bioscience (Bristol, UK).

### Hippocampal slice preparation

Immediately after decapitation, the brain was dissected and placed in ice-cold artificial cerebrospinal fluid (aCSF) containing (in mM): 124 NaCl, 30 NaHCO_3_, 10 glucose, 1.25 NaH_2_PO_4_, 3.5 KCl, 1 MgCl_2_, and 2 CaCl_2_. Horizontal hippocampal slices (400-µm thickness) were prepared using a Leica VT1200 vibratome (Leica; Deerfield, IL) and placed in an interface incubation chamber containing aCSF. The chamber was kept at 34°C during slicing and then returned to ambient room temperature for at least 2 hr. The slices were continuously bathed in aCSF bubbled with humidified carbogen gas (5% CO_2_/95% O_2_).

### fEPSP recordings

Acute hippocampal slices were transferred to a submerged recording chamber and continuously perfused at 2-3 ml/min with aCSF at 32°C. An extracellular borosilicate recording pipette filled with aCSF was placed in the stratum radiatum, and field excitatory postsynaptic potentials (fEPSPs) were evoked by electrical stimulation of the Schaffer collaterals (50-µs duration) using a bipolar concentric electrode (FHC Inc., Bowdoin, ME) connected to an isolated current stimulator (Digitimer Ltd., Welwyn Garden City, UK). Recordings were performed with the stimulus intensity set to elicit 30-40% of the maximal response (typically 30-50 µA), and individual synaptic responses were evoked at 0.1 Hz (every 10 s), unless otherwise stated. Paired-pulse facilitation was recorded with an interstimulus interval of 50 ms and the paired-pulse facilitation ratio (PPR) was calculated by dividing the fEPSP slope evoked by the second pulse by the fEPSP slope evoked by the first pulse. For recording the NMDA receptor–mediated component of the fEPSP, responses were evoked in the presence of 50 µM NBQX with 0.2 mM MgCl_2_ in the aCSF; at the end of the recording, the specificity of the signal was confirmed by applying 25 µM AP5. The acquired signal was amplified and filtered at 2 kHz (low-pass filter) using an extracellular amplifier (EXT-02F, NPI Electronic, Tamm, Germany). Data were collected and analyzed using a Digidata 1440A, Axoscope, and Clampfit (Molecular Devices, Can Jose, CA). Responses were quantified by determining the slope of the linear rising phase of the fEPSP (between 10% and 70% of the peak amplitude). The response was then normalized to the average baseline measured in the last 5 min prior to the start of the experiment. Time courses were created by averaging 6 fEPSPs collected during 1 min. For fEPSP responses evoked at 0.001 Hz, 3 single stimuli delivered at 0.1 Hz were applied and averaged at 15-min intervals. To induce LTP in the CA1 region, theta-burst stimulation (θ-burst; 10 bursts of 4 pulses at 100 Hz, delivered at 5 Hz) was applied twice with a 10-s interval. For subthreshold stimulation, the number of θ-bursts was determined in control slices at the beginning of each experiment and ranged from 5 to 8 bursts. Where indicated, the data were re-normalized to the average of the last 5 min prior to inducing LTP. The presence of LTP was determined by comparing the average of the fEPSP slope measured 55-60 minutes after θ-burst stimulation with the average of the last 5 min prior to stimulation. For group comparisons, time-matched 5-min averages were used, unless otherwise indicated. The rate of change of the fEPSPs was estimated by measuring the slope of the average time course, calculated using a linear regression with least-squares fitting during the times indicated in each figure.

### Patch-clamp recordings

After the acutely prepared slices recovered for least one hour, they were transferred to a submerged recording chamber perfused at 2-3 ml/min with aCSF at 32±1°C. The Schaffer collaterals were stimulated at a frequency of 0.1 Hz using a bipolar electrode, and the corresponding field potential was recorded. After a baseline recording of 10 min, DL-TBOA (50 μM) was added. During 40 min in DL-TBOA, the synaptic retuning and new steady state was induced and confirmed as above, after which the neurons in the CA1 layer (identified by their shape and localization in the CA1 pyramidal cell layer) proximal to the recording electrode were recorded using a Ag/AgCl electrode in a borosilicate glass pipette with a tip resistance of 4-5 MΩ filled with a solution containing (in mM): 110 K-gluconate, 10 KCl, 4 Mg-ATP, 10 Na_2_-phosphocreatine, 0.3 Na-GTP, 10 HEPES, and 0.2 EGTA (pH 7.2-7.4; 270-290 mOsm).

To isolate miniature EPSCs arising from spontaneously released synaptic vesicles, 1 µM tetrodotoxin was added to the perfusion solution to block action potentials. Access resistance was monitored throughout the recordings, and only stable recordings (<20% variation) were included in our analysis. Data were acquired using a Multiclamp 700B amplifier and Clampex 10.0 (Molecular Devices) and digitized using a Digidata 1440A (Molecular Devices). Traces were analyzed using the Mini Analysis Program (Synaptosoft Inc., Fort Lee, NJ).

### Enzyme-based glutamate sensor

The ceramic-based microelectrode array (MEA; CenMeT Service Center, Lexington, KY) consisted of two 50 μm × 150 μm platinum recording sites arranged in a row with 50 μm spacing between sites. One recording site was designed to be sensitive to glutamate and was coated with 2% glutamate oxidase (US Biological, G4001-01), 1% BSA (Sigma-Aldrich, A3059), and 0.125% glutaraldehyde (Sigma-Aldrich, G5882); the other site served as a sentinel detector and was coated only with BSA and glutaraldehyde. To prevent potential interference from reaching the MEA surface and to increase selectivity, the enzyme-coated MEAs were electroplated with the size-exclusion polymer m-phenylenediamine (mPD) by cycling between +0.2 V and +0.8 V at 50 mV/s for 22 min in a nitrogen-bubbled solution containing 10 mM mPD. Coated MEAs were allowed to dry for 48 h at room temperature in low humidity prior to *in vitro* calibration. The glutamate signal was obtained by subtracting the signal measured at the sentinel site from the signal measured at the enzyme-coated site.

The microelectrodes were calibrated each day prior use in slice recording. Constant potential amperometry was performed by applying a potential of +0.7 V relative to an Ag/AgCl reference electrode. Calibrations were performed in a stirred solution of 0.05 M phosphate-buffered saline (pH 7.4, 37°C). After a stable baseline signal was achieved, 250 µM ascorbic acid, 20 µM L-glutamate (3x), 2 µM dopamine (freshly prepared), and 8.8 µM H_2_O_2_ were added sequentially to the calibration beaker. Amperometric signals were acquired at 0.1 Hz using a FAST-16 mk-I electrochemical recording system (Quanteon LLC, Nicholasville, KY) and analyzed off-line using FAST Analysis software (Quanteon LLC).

### Propidium iodine (PI) exclusion

To measure cell viability, slices were prepared as described above; after recovering in the interface chamber, the slices were transferred to a submerged chamber and incubated with 50 µM DL-TBOA or 10 mM NaN_3_ for 1 h. The slices were then transferred to an incubation chamber containing propidium iodine (1 µg/ml) for 15 minutes, washed, and mounted in medium containing DAPI (Vector Laboratories, Burlingame, CA). The slices were imaged using an epifluorescence microscope, and both PI-positive cells and DAPI-stained nuclei were counted. The fraction of dead cells was calculated as the number of PI-positive cells divided by the number of DAPI-stained nuclei.

### Biotinylation of cell-surface proteins and Western blot

Hippocampal slices were pre-treated with aCSF alone, 50 µM DL-TBOA or 50 µM DL-TBOA with 10 µM MK-801 during 40 min and subsequently washed-out in ice-cold aCSF. Cell surface proteins were then biotinylated using 100 µg/µl sulfo-NHS-SS-Biotin (Pierce; A39258) for 30 min at 4°C with gentle shaking. After wash-out, free biotin was quenched in 100 mM glycine buffer (15 min, x2, 4°C). After wash-out, hippocampal biotinylated slices were lysed in ice-cold RIPA buffer containing protease inhibitors.

Protein concentration in the samples was determined using BCA assay (Pierce; 23227). Biotinylated proteins in 100 µg of total lysate were isolated by binding to NeurtrAvidine beads (Pierce; 100 µl) and eluted using SDS-PAGE sample buffer +5% β-mercaptoetanol. Total protein samples and eluted biotinylated fractions were analyzed by western blot. After primary antibody incubation (GluA2; anti-GluR2 ab20673, rabbit polyclonal, 1:100; Abcam, UK), the final detection of proteins of interest was based on a fluorescent secondary antibody detection (donkey anti-rabbit 1:20000) using the LICOR Odyssey infrared fluorescence system. Densitometric analysis were conducted using ImageJ (NIH, Bethesda, MD, USA).

## Supporting information

SuppMat

## Acknowledgments

We thank Carl Björkholm, Marta Gómez-Galàn, and members of the Lindskog lab for valuable input throughout this work.

## Competing interests

Authors declare that they have no competing interests.

The work was funded by:

Stiftelsen Olle Engkvist Byggmästare (ML)

Swedish Brain Foundation FO2018-0155 (ML)

Swedish Research Council Dr. Nr. 2017-02812 (ML)

Magnus Bergvall Foundation (ML)

Conacyt Post doc funding (E.V.-J.)

## Competing interests

Authors declare that they have no competing interests.

The work was funded by:

Stiftelsen Olle Engkvist Byggmästare (ML)

Swedish Brain Foundation FO2018-0155 (ML)

Swedish Research Council Dr. Nr. 2017-02812 (ML)

Brain and Behavior Research Foundation No 24571 (ML)

Magnus Bergvall Foundation (ML)

Conacyt Post doc funding (E.V.-J.)

## Data and materials availability

All data used in the analysis is available upon request from corresponding author.

## Notes

### Competing Interest Statement

The authors have declared no competing interest.

