## Supplementary material for "Impaired glutamate reuptake induces synaptic mistuning in rat hippocampal slices, that can be counteracted by ketamine": SuppMat

### Supplementary Figure 1-4

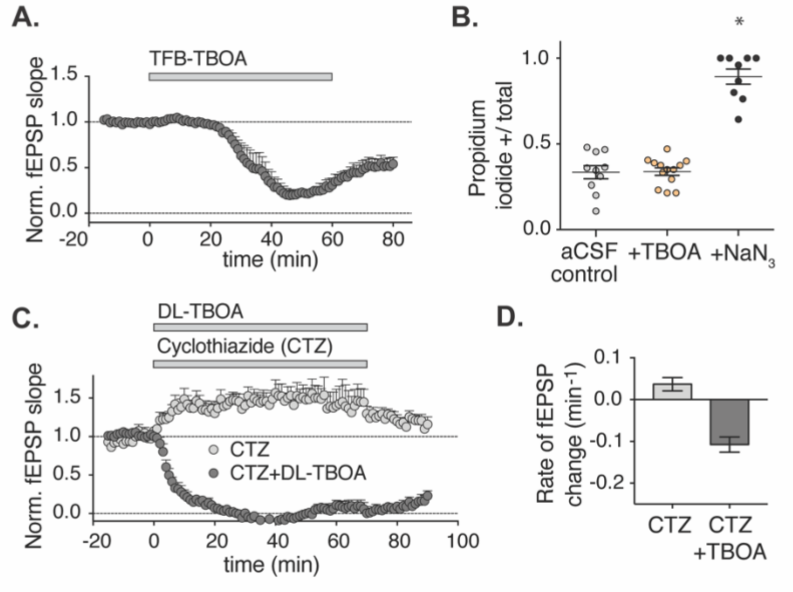


**Supplementary figure 1.**

Fig. S1. A) TFB-TBOA, a glutamate transporter blocker with higher specificity for isoforms expressed mainly in astrocytes, evoked the same re-tuning of the synapse as DL-TBOA. fEPSP slope over time plotted as average of 6 stimulation pulses per minute (200 nM; n=4). B) The effect of 60 min exposure to DL-TBOA (n=13), aCSF alone (n=10) or sodium azide (1mM, n=9) on cell viability determined by propidium iodine staining and presented as fraction of total number of cells that have taken up propidium iodine. C) fEPSP over time at application of DL-TBOA in the presence of 50μM cyclothiazide (n=6) and the effect of cyclothiazide alone (n=4). D) Rate of change of fEPSP slope at minute 1-5 after drug application. Error bars denote standard error of the mean (SEM) and *=p<0,05 ANOVA multiple comparison test.

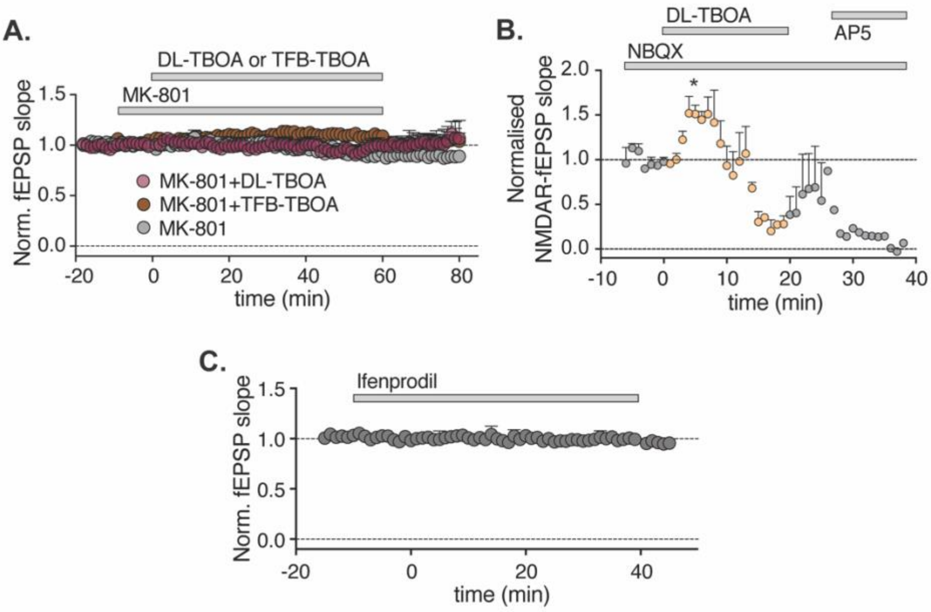


**Supplementary figure 2.**

Fig. S2. A) Time-course of fEPSP at application of 50μM DL-TBOA or 200nM TFB-TBOA in the presence of 10μM MK-801 (pre-incubation, 10 min; n=4 and 2 respectively), versus MK-801 alone (n=3). B) NMDA receptor-mediated component of the fEPSP over time (25 μM NBQX in 0.2 mM MgCl2 and fully inhibited by AP5) at application of 50 μM DL-TBOA and its'wash-out. C) fEPSP over time at application of 10 μM ifenprodil (n=2) *=p<0,05 Mann-Whitney U test vs baseline.

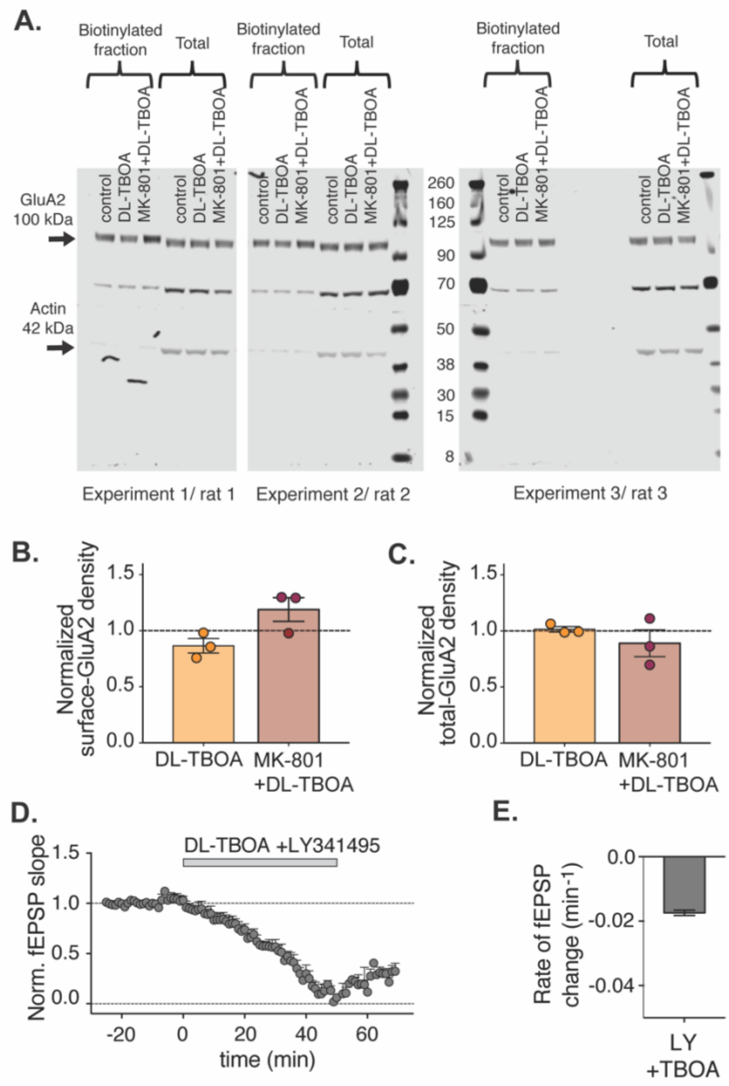


**Supplementary figure 3.**

Fig. S3. A) Surface levels of GluA2 after 40 min treatment with aCSF control, 50µM DL-TBOA or 10 µM MK-801 + 50 µM DL-TBOA were assessed using brain slice cell surface biotinylation procedure; equal amounts in µg of total lysates were analyzed in parallel to determine the total GluA2 expression levels. B) Densitometric analysis of cell-surface GluA2 (biotinylated fraction) normalized to control (n=3). C) Densitometric analysis of GluA2 in total lysate normalized to control (n=3). D) Time course of normalized fEPSP slope; where indicated, 50 µM DL-TBOA combined with 1 µM LY341495 a Group II mGlu receptor inhibitor was added (n=5 slices). E) Rate of change in the fEPSP response over a 35 min period after DL-TBOA + LY341495 addition and measured as the slope of a first order linear model fitted to the fEPSP time-response.

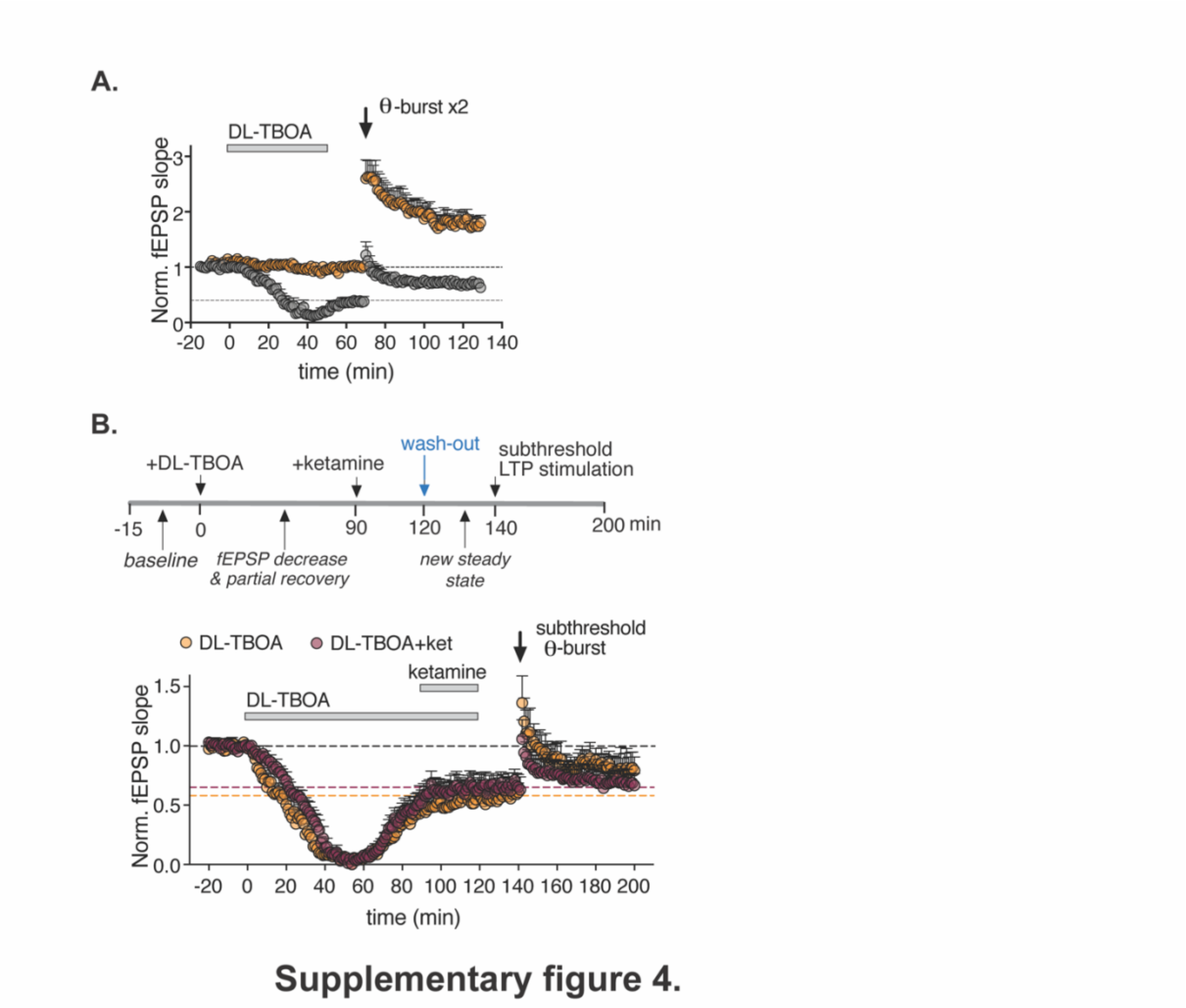


Fig. S4. A) fEPSP over time (full time-course from Fig. 4F) normalized to initial fEPSP baseline. 20 minutes after DL-TBOA wash-out, at the new steady state, Schaffer collaterals were stimulated with a standard LTP inducing theta burst applied at 60 minutes, n=5 for control, 4 for DL-TBOA treated. B) fEPSP over time (full time-course from Fig. 5E) normalized to initial fEPSP baseline, 90 minutes after DL-TBOA application, n=5 for ketamine + DL-TBOA treated slices, n=11 for DL-TBOA treated slices. Error bars denote standard error of the mean (SEM).
